# Increased GABA transmission to GnRH neurons after intrahippocampal kainic acid injection in mice is sex-specific and associated with estrous cycle disruption

**DOI:** 10.1101/2022.05.20.492873

**Authors:** Robbie J. Ingram, Leanna K. Leverton, Victoria C. Daniels, Jiang Li, Catherine A. Christian-Hinman

## Abstract

Patients with epilepsy develop reproductive endocrine comorbidities at a rate higher than that of the general population. Clinical studies have identified disrupted luteinizing hormone (LH) release patterns in patients of both sexes, suggesting potential epilepsy-associated changes in hypothalamic gonadotropin-releasing hormone (GnRH) neuron function. In previous work, we found that GnRH neuron firing is increased in diestrous females and males in the intrahippocampal kainic acid (IHKA) mouse model of temporal lobe epilepsy. Notably, GABA_A_ receptor activation is depolarizing in adult GnRH neurons. Therefore, here we tested the hypothesis that increased GnRH neuron firing in IHKA mice is associated with increased GABAergic drive to GnRH neurons. When ionotropic glutamate receptors (iGluRs) were blocked to isolate GABAergic postsynaptic currents (PSCs), no differences in PSC frequency were seen between GnRH neurons from control and IHKA diestrous females. In the absence of iGluR blockade, however, GABA PSC frequency was increased in GnRH neurons from IHKA females with disrupted estrous cycles, but not saline-injected controls nor IHKA females without estrous cycle disruption. GABA PSC amplitude was also increased in IHKA females with disrupted estrous cycles. These findings suggest the presence of an iGluR-dependent increase in feed-forward GABAergic transmission to GnRH neurons specific to IHKA females with comorbid cycle disruption. In males, GABA PSC frequency and amplitude were unchanged but PSC duration was reduced. Together, these findings suggest that increased GABA transmission helps drive elevated firing in IHKA females on diestrus and indicate the presence of a sex-specific hypothalamic mechanism underlying reproductive endocrine dysfunction in IHKA mice.

**HIGHLIGHTS:**

- Increased GABA transmission to GnRH neurons in IHKA mouse model of epilepsy
- Increased GABA transmission is dependent on upstream glutamate signaling
- Increased GABA transmission only seen in females with disrupted estrous cycles
- Potential sex-specific mechanism for reproductive endocrine dysfunction in epilepsy

## INTRODUCTION

Epilepsy is the fourth leading neurologic disorder in the United States and affects the lives of approximately 3 million Americans (England et al., 2012; Zack and Kobau, 2017). Temporal lobe epilepsy (TLE) is the most common form of focal epilepsy in patients of reproductive age (Engel, 2001). Importantly, people with TLE exhibit multiple forms of reproductive endocrine dysfunction at rates higher than the general population (Bauer and Cooper-Mahkorn, 2008; Bilo et al., 2001; Herzog et al., 1986a, 1986b; Klein et al., 2001). While certain antiseizure drugs can elicit reproductive endocrine dysfunction in people with epilepsy (Isojärvi et al., 2005), it should be noted that these comorbidities can also arise in the absence of pharmacological antiseizure treatment (Bilo et al., 1988; Herzog et al., 1986a, 1986b). Additionally, clinical studies have found that seizure focus location plays a role in determining the likelihood of different reproductive endocrine comorbidities (Bauer et al., 2004; Herzog, 1993; Kalinin and Zheleznova, 2007). Taken together, these studies suggest that seizure activity itself can increase susceptibility to reproductive endocrine comorbidities.

In females, these comorbidities can take the form of polycystic ovary syndrome (Bauer et al., 2000; Herzog, 2008), menstrual cycle disruption (Herzog, 2008; Herzog et al., 2003; Zhou et al., 2012), and/or hormonal disruption (Herzog et al., 2003). In males, comorbidities such as semen abnormalities (Herzog et al., 1986b) and low serum testosterone (Talbot et al., 2008) are observed. Notably, disrupted patterns of luteinizing hormone (LH) release from the pituitary gland have been documented in both male and female TLE patients (Drislane et al., 1994; Herzog et al., 1990, 1986a; Quigg et al., 2002).

In the mammalian brain, gonadotropin-releasing hormone (GnRH) neurons serve as the final common pathway for the neural control of reproductive endocrine function. The pulsatile release of GnRH is the primary driver of properly timed pituitary LH pulses (Belchetz et al., 1978; Haisenleder et al., 1991; Wildt et al., 1981). Due to the tightly integrated nature of GnRH and LH release, GnRH neurons are likely mediators of epilepsy-associated reproductive endocrine dysfunction. Recent work documented that GnRH neuron firing activity and intrinsic excitability are altered in the intrahippocampal kainic acid (IHKA) mouse model of TLE (Li et al., 2018). GABAergic fast synaptic transmission provides a primary regulatory input to GnRH neurons (Christian and Moenter, 2008, 2007), with relatively little direct input from glutamatergic afferents (Christian et al., 2009; Shimshek et al., 2006; Suter, 2004). Notably, GnRH neurons maintain depolarizing, rather than hyperpolarizing, responses to GABA_A_ receptor (GABA_A_R) activation in adulthood (DeFazio et al., 2002; Herbison and Moenter, 2011; Moenter and DeFazio, 2005; Taylor-Burds et al., 2015). Therefore, we hypothesized that epilepsy-associated increases in GnRH neuron firing are driven, at least in part, by increased afferent GABAergic drive.

To determine whether the epilepsy-associated changes in GnRH neuron firing activity in the IHKA model are associated with altered GABAergic transmission, we recorded GABA_A_R-mediated postsynaptic currents (PSCs) in GnRH neurons in slices of hypothalamus from IHKA mice and saline-injected controls. All recordings were performed at 2-3 months after injection, a time point by which a large majority of IHKA mice have been documented to display chronic spontaneous seizures (Cutia et al., 2022; Li et al., 2020; Lisgaras and Scharfman, 2022) and the majority of IHKA females exhibit disrupted estrous cycles (Cutia et al., 2022; Li et al., 2017). In light of the largely depolarizing effects of GABA_A_R activation in GnRH neurons, which could potentially promote increased firing activity, we recorded GABAergic transmission to GnRH neurons from males and diestrous females, as increased GnRH neuron firing was previously observed in IHKA mice from these groups (Li et al., 2018).

## MATERIALS AND METHODS

### Animals

GnRH-tdTomato transgenic mice on the C57BL/6J background were bred by crossing GnRH-Cre^+^ females ((Yoon et al., 2005); The Jackson Laboratory #021207) and Ai9 males ((Madisen et al., 2010); The Jackson Laboratory #007909). Mice were housed with a 14:10 h light:dark cycle (7:00 PM lights-off) to promote estrous cyclicity and breeding activity (Fox et al., 2006). Before IHKA injection, housing consisted of a standard environment with up to five mice per cage. Mice within each litter were randomly assigned to the saline or KA group in order to minimize potential litter effects. At the time of IHKA injection, female litters were separated into smaller groups depending on whether their days of diestrus aligned, allowing for injection on the same day and housing in the same cage after injection. Pups were genotyped via PCR of DNA extracted from tail clips collected before postnatal day (P)21. The following four primer sequences were used, as suggested by The Jackson Laboratory: (1) transgene reverse CGG ACA GAA GCA TTT TCC AG; (2) transgene forward ACA GGT GTC TGT CCC ATG TCT; (3) internal positive control forward CAA ATG TTG CTT GTC TGG TG; (4) internal positive control reverse GTC AGT CGA GTG CAC AGT TT. All animal procedures were approved by the Institutional Animal Care and Use Committee of the University of Illinois Urbana-Champaign (protocols 17183 and 20155).

### Estrous cycle monitoring

Prior to injection, regular estrous cyclicity in females was confirmed via daily vaginal smears obtained between 10:00 AM and 12:00 PM from P42 onwards (Pantier et al., 2019). 20 µl of sterile PBS was inserted into the vaginal cavity, removed, and placed on a glass slide for examination via brightfield microscopy. Smears were classified as one of the 5 following stages based on the primary cell type(s) present: (1) proestrus: nucleated epithelial cells; (2) estrus: cornified epithelial cells; (3) metestrus: both cornified epithelial cells and leukocytes; (4) diestrus I: leukocytes; (5) diestrus II: few or no cells present. Estrous cycle stage was recorded daily for at least 14 days and the average cycle length for each mouse was calculated. The mouse estrous cycle is typically 4-5 days in length (Byers et al., 2012). Only mice determined to have regular estrous cycles underwent an intrahippocampal injection of either saline or KA. After one month of undisturbed rest, daily estrous cycle monitoring resumed. Cycle period phenotype was assessed from the data recorded from 42 days post-injection onwards, which was previously identified as a time by which cycle disruption presents in IHKA mice (Li et al., 2017). To provide a buffer against false positives from minor disruptions in cyclicity, only mice with cycles ≥ 7 days in length were categorized as having a disrupted cycle and classified as “KA-long”, whereas IHKA mice with regular cycles (average length 4-6 days) were classified as “KA-regular” (Li et al., 2018).

### Intrahippocampal injections

Stereotaxic injections were performed in male and diestrous female mice older than P56 under 2-3% oxygen-vaporized isoflurane anesthesia (Clipper Distributing Company). For analgesia, carprofen (5 mg/kg, Zoetis) was administered via subcutaneous injection at the beginning of the surgery. KA (Tocris Bioscience; 50 nl of 20 mM prepared in 0.9% sterile saline) was injected into the right dorsal hippocampal CA1 region (coordinates: 2.0 mm posterior and 1.5 mm lateral to bregma; 1.4 mm ventral to the cortical surface). Control mice were injected with an equivalent volume of sterile saline in an identical manner. After completion of the injection, the incision was closed with sutures. Following closure of the wound, anesthetic 2.5% lidocaine + 2.5% prilocaine cream (Hi-Tech Pharmacal) and Neosporin antibiotic gel (Johnson and Johnson) were applied to the affected area.

### Video monitoring of acute behavioral seizures

Immediately after intrahippocampal injection, mice were placed in a warmed and transparent recovery chamber. All KA-injected mice were video recorded for 3-5 hours using Logitech webcams and a Windows PC. Videos were then screened for acute behavioral seizures of Racine stage 3 (forelimb clonus) and higher (rearing and falling). Seizures of lower Racine stages were not noted, as they could not be reliably distinguished from normal mouse behavior in the video recordings.

### Brain slice preparation

Acute brain slices were prepared 2-3 months after injection surgery following procedures described previously (Li et al., 2018). For females, all slice experiments were done during diestrus. Briefly, mice were euthanized via decapitation between 10:00 and 11:00 AM. Coronal brain slices of 300-µm thickness were prepared using a Leica VT1200S vibrating blade microtome (Leica Biosystems). During slicing, brain slices were submerged in an oxygenated (95% O_2_, 5% CO_2_) ice-cold sucrose solution (2.5 mM KCl, 1.25 mM NaH_2_PO_4_, 10 mM MgSO_4_, 0.5 mM CaCl_2_, 11 mM glucose, and 234 mM sucrose). Brain slices were transferred to an oxygenated artificial cerebrospinal fluid (ACSF) solution (2.5 mM KCl, 10 mM glucose, 126 mM NaCl, 1.25 mM NaH_2_PO_4_, 1 mM MgSO_4_, 0.5 mM CaCl_2_, and 26 mM NaHCO_3_; osmolarity ≈ 298 mOsm) for 30 min at 32°C, then transferred to room temperature for at least 15 min before recording. For recording, each slice was individually placed in the recording chamber of an upright BX51W1 microscope (Olympus America) and submerged with oxygenated ACSF warmed to 32°C using an inline heater (Warner Instruments) and circulated through the slice chamber at a flow rate of 2-3 ml/min.

### Voltage-clamp recordings

The experimenter was blinded to the injection group of each mouse during recording and analysis. All recordings were performed between 1:00-4:00 PM and targeted tdTomato-expressing GnRH neurons in coronal brain slices containing the medial septum (MS) and/or preoptic area (POA). All recorded cells were ipsilateral to the intrahippocampal injection. Soma position was noted with the use of a mouse brain atlas ((Paxinos and Franklin, 2019); corresponding plates: MS = 23-25 and POA = 25-28). Borosilicate glass pipettes (Sutter Instruments) were pulled with a Model P-1000 Flaming/Brown Micropipette Puller (Sutter Instruments), with a target tip resistance of 2-3 MΩ when filled with a CsCl-based (135 mM CsCl, 10 mM HEPES, 10 mM EGTA, 2 mM MgCl_2_, and 5 mM QX-314; osmolarity ≈ 290 mOsm) internal pipette solution (Christian et al., 2013; Courtney and Christian, 2018). The pipette was positioned with an MPC-2000 micromanipulator (Sutter Instruments), and data recorded via an Axon MultiClamp 700B amplifier (Molecular Devices) paired with an Axon Digidata 1550 Digitizer (Molecular Devices). Recordings were performed using Clampex 10.7 software (Molecular Devices). Imaging through a Retiga R1 CCD camera (Teledyne Photometrics) was controlled with µManager 1.4 software (Edelstein et al., 2010).

Once a GΩ seal was obtained and whole-cell configuration achieved, the cell was allowed to stabilize for >120 s prior to recording. After this stabilization period, passive membrane properties were assessed using an access test with a hyperpolarizing voltage step from -60 mV to -65 mV (mean of 20 repeats, 20 ms duration, sampled at 50 kHz with gain set at 5, Bessel filtered at 20 kHz) before and after each 2-min gap-free PSC recording. Membrane potential was clamped at -60 mV at all times, except during the hyperpolarizing step within the access test. Only recordings with a stable R_in_, stable R_s_ < 20 MΩ, stable C_m_ greater than 7.5 pF, and stable I_hold_ between 0 and -100 pA were included in analyses. An analysis of these properties showed that GnRH neurons from KA-long females exhibited an increased C_m_ (25.98 ± 2.61 pF) when compared to neurons from saline-injected females (17.98 ± 1.64 pF; *p* = 0.03, two-way ANOVA/Tukey’s post-hoc), but there were no other differences in passive cell properties. Gap-free recordings were also sampled at 50 kHz, with the gain set to 10 and filter set to 4 kHz. Two 2-min bins of PSC recordings were used in analyses for each cell. Where indicated, 1 mM kynurenic acid (Sigma K3375) was added to the circulating ACSF bath solution. Recordings were analyzed with StimFit 0.15 (Guzman et al., 2014).

### Hippocampal histology

After hypothalamic slice preparation, the remaining portion of the brain containing the hippocampus was collected from the vibratome chuck and bulk-fixed in 4% paraformaldehyde for 24 h at 4°C. It was then stored in 30% sucrose solution with 0.5% sodium azide at 4°C until preparation of 40 µm-thick coronal sections using a freezing microtome (SM 2010R, Leica Biosystems). Four to eight sections per mouse through the dorsal hippocampal region were used to evaluate hippocampal sclerosis by cresyl violet staining or gliosis by glial fibrillary acidic protein (GFAP) staining.

For cresyl violet staining, sections were mounted on charged glass slides and stained with cresyl violet (Sigma C5042) for 12 min at room temperature (∼22°C). They were then dehydrated with a series of graded ethanol solutions (70-100%), cleaned in xylene, and coverslipped with Permount. For GFAP immunostaining, floating sections were incubated in an anti-GFAP mouse monoclonal primary antibody (1:1000, Sigma G3893) on a shaker for 48 h at 4°C, followed by incubation in Fluorescein-conjugated horse anti-mouse secondary antibody (1:1000, Vector Laboratories Fl-2000) on a shaker for 2 h at room temperature. After staining, all sections were mounted on charged glass slides and coverslipped using Vectashield Vibrance Antifade Mounting Medium with DAPI (Vector Laboratories H-1800). Image acquisition was performed using an Olympus BX43 microscope with an Infinity 3-6UR camera and Infinity Capture software (Teledyne Lumenera).

### Statistical analyses

Statistical comparisons were made using R software with custom code. Comparisons between two groups were made via *t* tests or Mann-Whitney tests, depending on the normality of the data. For two-group comparisons, normality was assessed via Shapiro-Wilk tests and homogeneity of variance was assessed via Levene’s tests. Comparisons between three groups were made via one-way ANOVA or Kruskal-Wallis tests, depending on the normality of the residuals and the homogeneity of variance between groups. Tukey’s and Dunn’s post-hoc tests were used for pairwise comparisons of significant ANOVA and Kruskal-Wallis results, respectively. Normality of residuals was assessed with Shapiro-Wilk tests, and homogeneity of variance was assessed with Levene’s tests. All results are reported as means ± SEM. Cumulative probability distributions containing a maximum of 100 randomly sampled PSCs (events) per cell were compared using Kolmogorov-Smirnov (K-S) tests (Christian et al., 2013; Christian and Moenter, 2007; Sullivan et al., 2003). Cells that did not have 100 total events had all available events included. Except for K-S tests between 3 groups, statistical significance for all analyses was set at *p* < 0.05. For K-S tests between 3 groups, the threshold for significance was set at a corrected p-value of *p* < 0.017 using the Bonferroni method. On average, 2 cells per mouse (maximum 3 per mouse) were recorded, and in the large majority of cases only one cell per slice was recorded.

Outliers were defined as values more than 2 standard deviations away from the group mean. According to this criterion, two outlier cells were removed, one from the KA-regular group in the kyn condition and one from the saline group in males. A separate analysis showed no effect of region (MS vs. POA) on PSC frequency in either females (*p* = 0.73, Kruskal-Wallis) or males (*p* = 0.81, one-way ANOVA). Therefore, the data from both regions were combined for both sexes.

## RESULTS

### Verification of successful intrahippocampal kainic acid injection

With the exception of estrous cycle monitoring, the same overall experimental procedure was used for both males and females (**Fig. 1**). To verify successful IHKA injections, we employed a three-layer screening procedure as done previously (Li et al., 2018). First, videos of mice recorded immediately after injection were screened for the occurrence of 2 or more behavioral seizures. Of 32 KA-injected mice, 26 displayed 2 or more seizures in this time frame. Next, hippocampal tissue from those mice that did not display at least 2 acute seizures after injection was then evaluated histopathologically for the presence of sclerosis and gliosis via cresyl violet (Nissl) or GFAP staining, respectively (**Fig. 2**). Of the 6 KA-injected mice not displaying 2 or more seizures post-injection (all females), no mice showed clear hippocampal sclerosis, but all showed clear signs of gliosis in the area of injection as indicated by GFAP staining (**Fig. 2B**). Therefore, all 32 KA-injected mice passed this screening procedure. It should be noted that we recently demonstrated that in a parallel cohort of mice injected identically to those described here, 100% developed spontaneous recurrent seizures by 1 month after injection, as recorded in hippocampal depth electroencephalography recordings (Cutia et al., 2022). This finding indicates a high rate of effective epilepsy induction in this model in our hands and provides strong confidence that the 32 KA-injected mice in the present work also developed chronic epilepsy by 2 months after injection.

**Figure 1:**
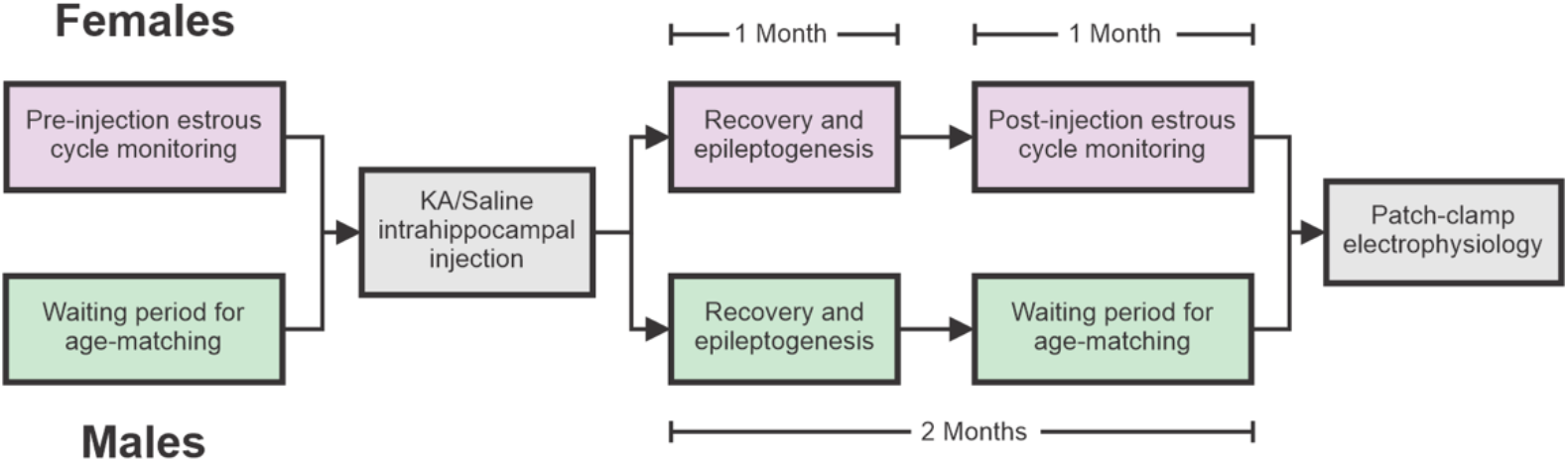
Flowchart illustrating similarities and differences between the experimental paradigms used for male and female mice. Note that all intrahippocampal injections and patch clamp electrophysiology experiments in females were performed using diestrous mice.

**Figure 2:**
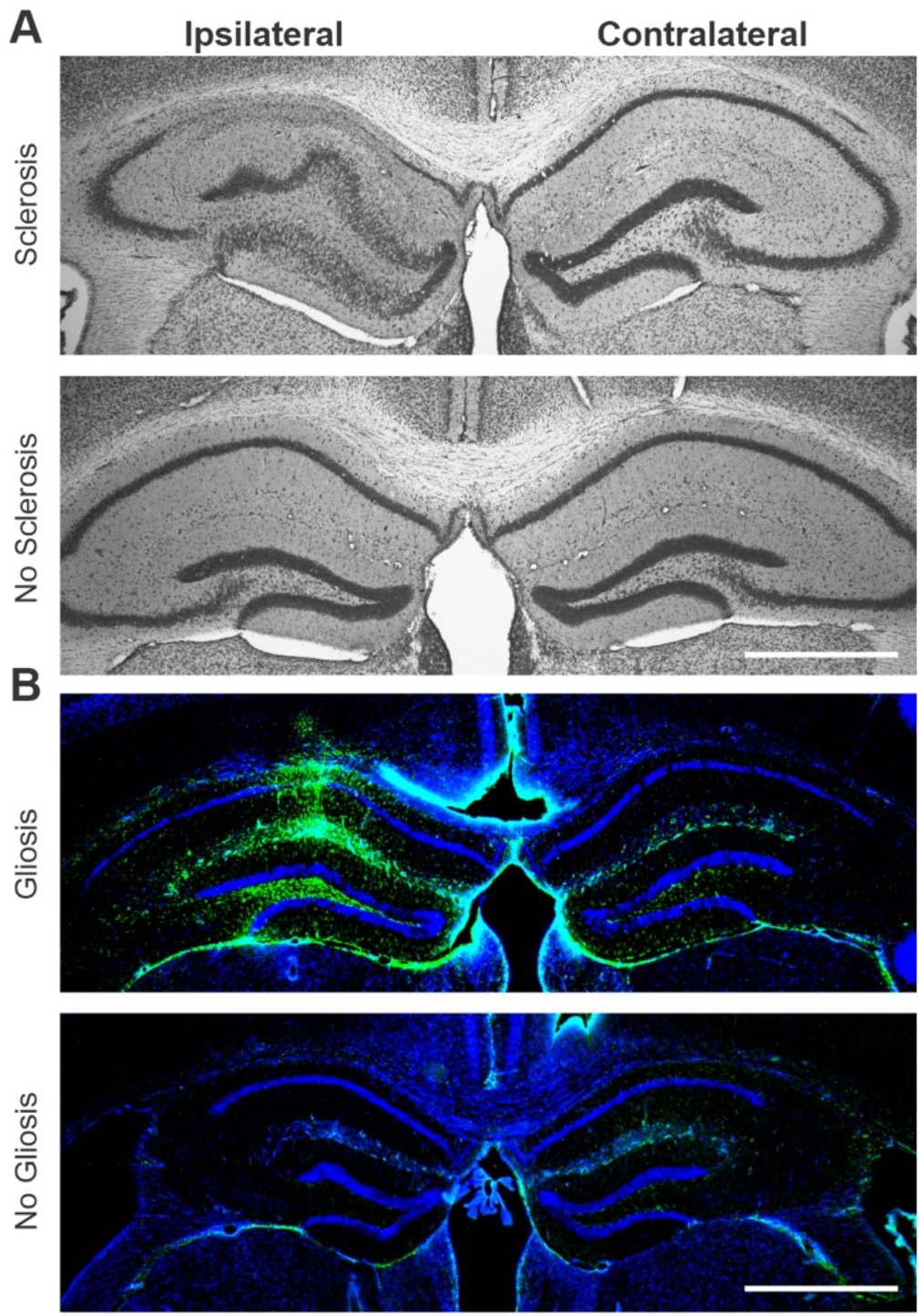
Histological verification of successful IHKA injection targeting. **A**, Example cresyl violet-stained sections showing presence (top) or absence (bottom) of sclerosis in ipsilateral hippocampus. **B**, Example GFAP (green) and DAPI (blue) staining showing presence (top) and absence (bottom) of gliosis in sections from IHKA mice. The example section exhibiting no gliosis is provided for visual comparison; other sections from the same mouse did exhibit signs of gliosis, and this mouse was thus included in the final data analyses. Note that gliosis was only examined in sections from IHKA mice that did not show signs of sclerosis in cresyl violet staining after screening for video-recorded seizures. Scale bars = 1 mm.

### Increased GABA sPSC amplitude in GnRH neurons from diestrous IHKA females in the presence of kynurenic acid

To identify potential changes in GABAergic transmission to GnRH neurons in diestrous females, we recorded GABA_A_R-mediated PSCs in the presence of 1 mM kynurenic acid to block ionotropic glutamate receptors (iGluRs) (**Fig 3**). In this condition, there were no differences in PSC frequency between saline (n= 10 cells, 7 mice), KA-long (n = 10 cells, 5 mice), and KA-regular (n= 11 cells, 7 mice) groups (F = 0.20, *p* = 0.82, one-way ANOVA) (**Fig. 3A-B**). An analysis of the cumulative probability distributions for GABA PSC amplitude showed a difference between KA-long (n = 918 events) and saline (n = 796 events, *p* < 6.65 × 10^−5^) and a difference between KA-regular (n = 960 events) and saline (*p* = 0.00203), with a shift towards larger amplitudes in the KA groups (**Fig 3C-D**). There was no difference in amplitude between KA-long and KA-regular groups (*p* = 0.19, K-S tests). An analysis of GABA PSC half width, a measure of duration, showed a slight difference between KA-regular and both KA-long (*p* = 3.96 × 10^−6^) and saline (*p* < 0.0055) (**Fig. 3E**). Sample sizes for analyses of PSC half width were identical to those used for amplitude. Overall, these findings indicate an increase in GABA sPSC amplitude in GnRH neurons from KA-long females, with differences in GABA sPSC half width having a less clear trend.

**Figure 3:**
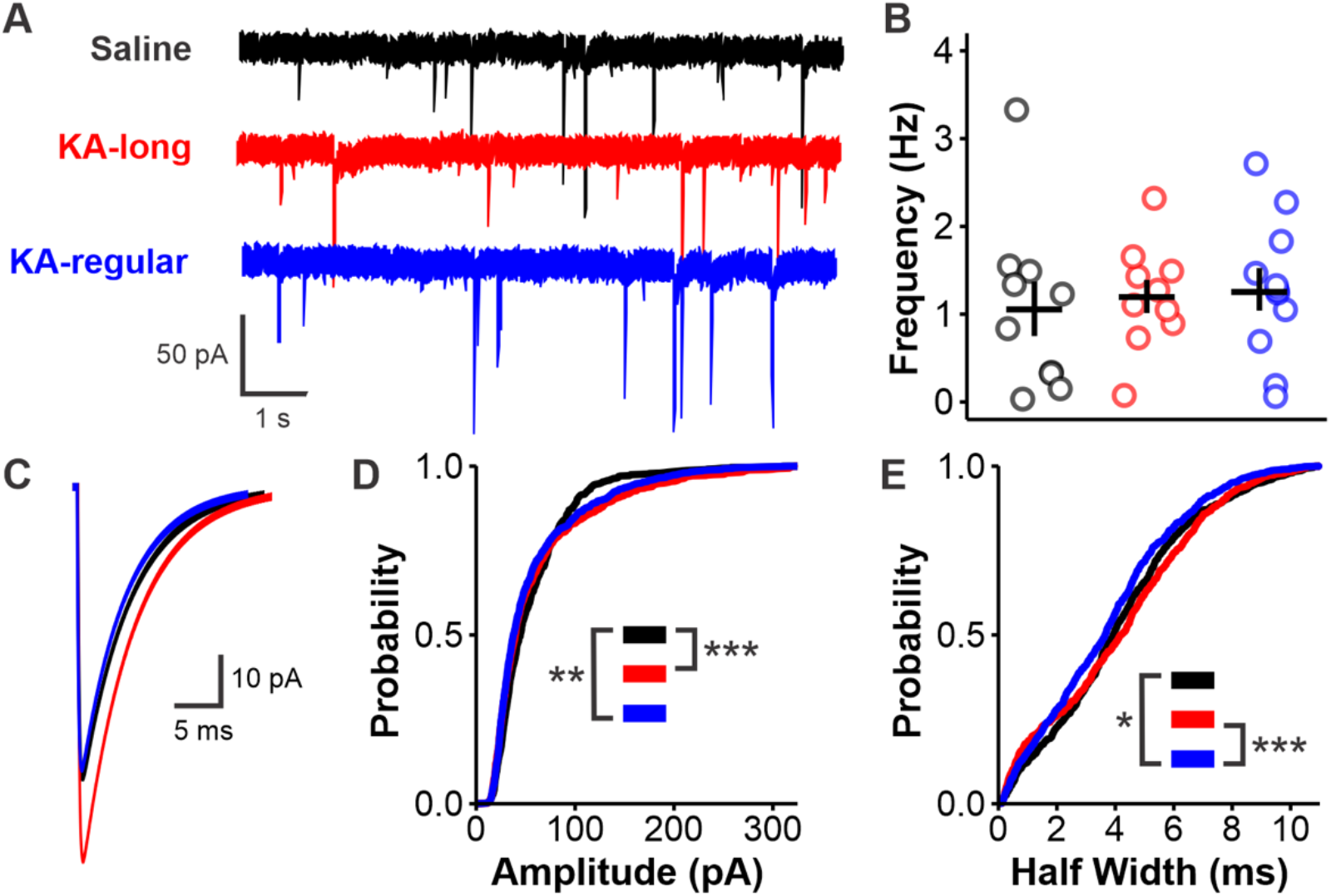
Increased GABA sPSC amplitude in GnRH neurons from diestrous female IHKA mice in the presence of iGluR blockade. **A**, Example traces. **B**, Scatter plot showing individual cell values (circles) and mean ± SEM for GABA PSC frequency in GnRH neurons from saline (black), KA-long (red), and KA-regular (blue) mice. **C**, Average PSC traces from all events obtained in each group. **D-E**, Cumulative probability distributions for PSC amplitude (**D**) and half width (**E**); \**p* < 0.017, \*\**p* < 0.003, \*\*\**p* < 0.0003 by Kolmogorov-Smirnov (K-S) tests corrected for multiple comparisons with the Bonferroni method.

### Restoring circuit integrity reveals iGluR-dependent increase in GABAergic transmission to GnRH neurons from KA-long female mice

The prior experiments that identified changes in GnRH neuron firing activity in IHKA mice were conducted without any receptor antagonists added to the ACSF bath (Li et al., 2018). Therefore, we were concerned that circuit integrity was being compromised by the addition of kynurenic acid, potentially masking glutamate-dependent increases in GABA transmission (i.e., via excitation of presynaptic GABAergic neurons). Recent work has highlighted the contribution of glutamatergic transmission to GABAergic neurons upstream of GnRH neurons (Wang et al., 2018), further supporting the notion of a potential iGluR-dependent increase in GABAergic transmission in IHKA mice.

Previous recordings of iGluR-mediated PSCs in GnRH neurons, as detected in somatic whole-cell recordings, have shown that the frequency of these currents is typically much lower than that of GABAergic PSCs (Christian et al., 2009; Liu et al., 2017; Suter, 2004). Furthermore, the kinetics of the iGluR-mediated PSCs are such that they can be distinguished from GABA_A_R-mediated PSCs by virtue of smaller amplitude values (Adams et al., 2018; Liu et al., 2017). Therefore, we tested whether the same recording configuration of a CsCl-based isotonic chloride pipette solution at V_hold_ = -60 mV would permit detection of GABAergic PSCs in GnRH neurons without any blockers in the bath, thus maintaining circuit integrity in the context of a coronal brain slice. Comparison of PSC amplitudes from recordings made with and without kynurenic acid present revealed a subpopulation of kynurenic acid-sensitive PSCs with amplitude values ranging from 0-15 pA (**Fig. 4A**). To account for this small but distinct population, a minimum amplitude cutoff of 15 pA to isolate putative GABAergic PSCs was applied to all recordings described below.

**Figure 4:**
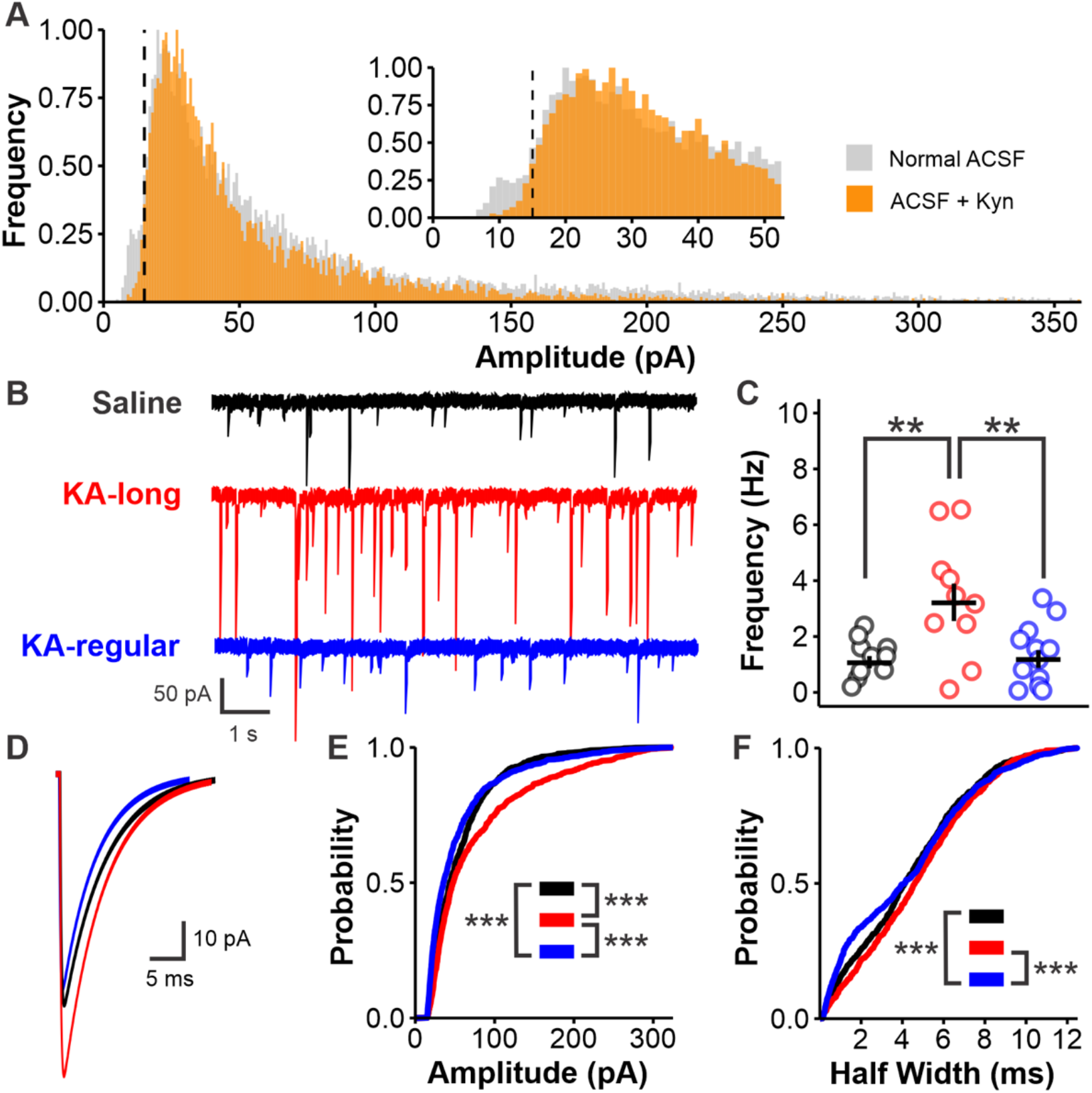
GABAergic transmission to GnRH neurons in female IHKA mice is increased in an iGluR-dependent manner. **A**, Histogram showing normalized distributions of unitary GABA PSC amplitude values from all recordings made in the presence (orange) or absence (grey) of 1 mM kynurenic acid. Dotted line indicates 15 pA minimum cutoff for putative GABAergic currents. **B**, Example traces recorded in control ACSF. **C**, Scatter plot showing individual cell values (circles) and mean ± SEM for GABA PSC frequency in GnRH neurons from saline (black), KA-long (red), and KA-regular (blue) mice; \*\**p* < 0.01 by Kruskal-Wallis with Dunn’s *post hoc* tests. **D**, Average PSC traces from all events obtained in each group. **E-F**, Cumulative probability distributions for PSC amplitude (**E**) and half width (**F**); \*\*\**p* < 0.0003 by Kolmogorov-Smirnov (K-S) tests corrected for multiple comparisons with the Bonferroni method.

In the absence of kynurenic acid, PSC frequency was increased in GnRH neurons from KA-long mice (n = 10 cells, 7 mice) compared with both control (n = 10 cells, 8 mice, *p* = 0.0091) and KA-regular groups (n = 12 cells, 7 mice, *p* = 0.0083; Kruskal-Wallis/Dunn’s) (**Fig. 4B-C**). This increase was specific to the KA-long group, as PSC frequency was not different between KA-regular and control groups (*p* = 0.47). An analysis of the cumulative probability distributions of GABA PSC amplitudes showed differences between all three groups, with KA-long (n = 918 events) shifted towards larger amplitudes in comparison to both saline (n = 948 events, *p* < 7.8 × 10^−9^) and KA-regular (n = 1120 events, *p* = 9.99 × 10^−14^) (**Fig 4D-E**). By contrast, the KA-regular group exhibited a slight shift towards smaller amplitudes in comparison to saline (*p* = 1.08 × 10^−6^, K-S tests). An analysis of the cumulative probability distributions of GABA PSC half width showed a difference between KA-regular and saline (*p* = 0.00016) as well as KA-long (*p* = 1.21 × 10^−9^), but there was no difference between saline and KA-long (*p* = 0.03064) (**Fig. 4F**). Sample sizes for analyses of PSC half width were identical to those used for amplitude. Overall, these findings suggest the presence of an iGluR-dependent increase in GABAergic transmission to GnRH neurons from KA-long females.

### GABAergic PSCs in GnRH neurons from IHKA male mice exhibit reduced duration, but no changes in frequency or amplitude

In males, there was no difference in mean GABA PSC frequency between KA (n = 14 cells, 6 mice) and saline (n = 15 cells, 12 mice) groups recorded in the absence of kynurenic acid (*p* = 0.6, t-test) (**Fig 5A-B**). An analysis of the cumulative probability distribution of GABA PSC amplitude showed no significant difference between KA (n = 1317 events) and saline groups (n = 1560 events, *p* = 0.28, K-S test) (**Fig. 5C-D**). A similar analysis of PSC half width, however, showed a reduction in duration in KA compared with saline (*p* = 1.0 × 10^−9^) (**Fig. 5E**), indicating potential postsynaptic changes in GABA_A_R subunit composition in GnRH neurons. Sample sizes for analyses of PSC half width were again identical to those used for amplitude. It should also be noted that a comparison between saline-injected males and females showed no baseline sex difference in GABA PSC frequency (*p* = 0.13, t-test). Overall, these findings indicate no major changes in GABAergic transmission to GnRH neurons from IHKA males.

**Figure 5:**
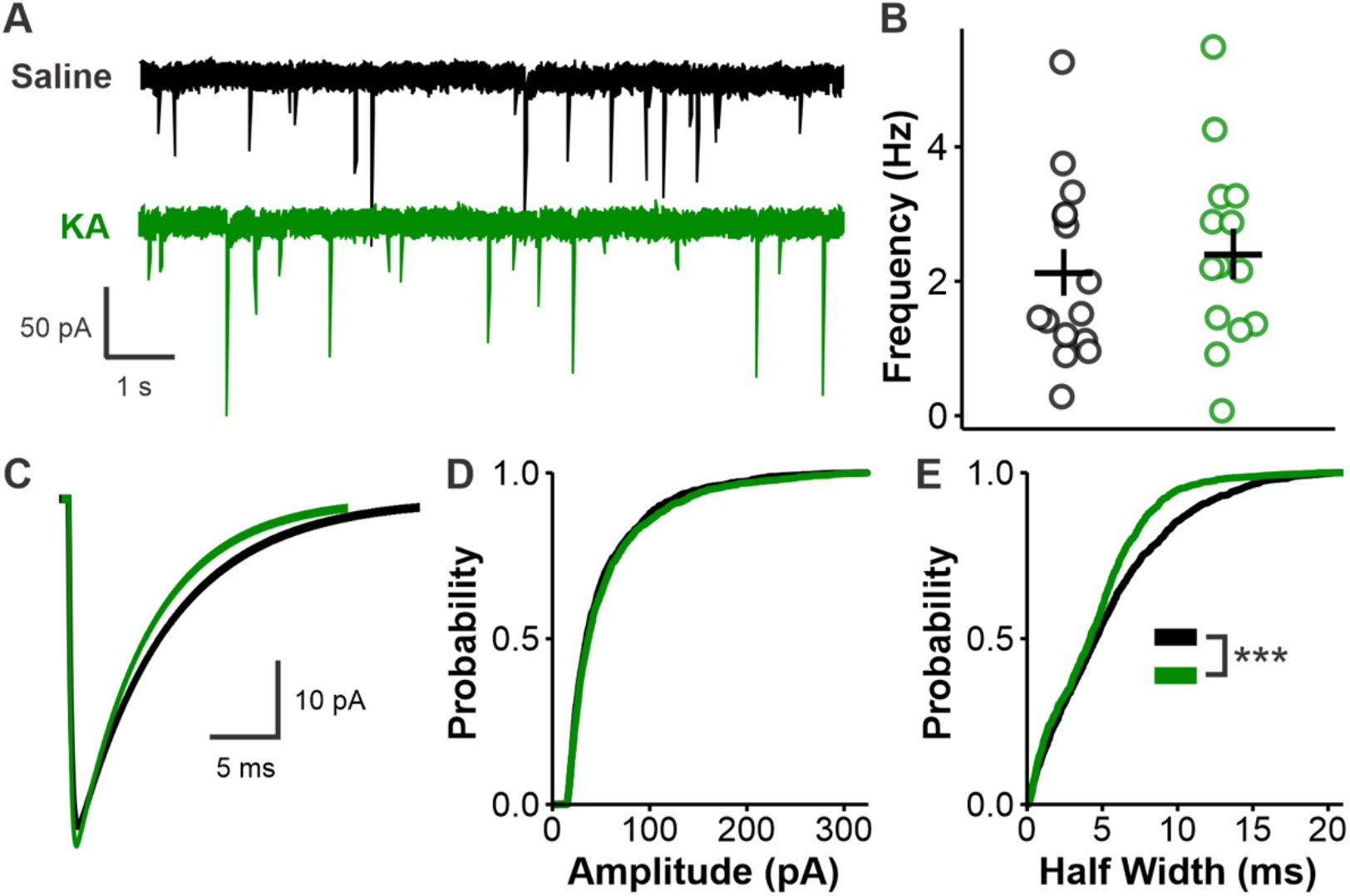
Reduced GABAergic PSC duration but no changes in frequency or amplitude in male IHKA mice. **A**, Example traces. **B**, Scatter plot showing individual cell values (circles) and mean ± SEM for GABA PSC frequency in GnRH neurons from saline (black) and KA (green) mice. **C**, Average PSC traces from all events obtained in each group. **D-E**, Cumulative probability distributions for PSC amplitude (**D**) and half width (**E**); \*\*\**p* < 0.001 by Kolmogorov-Smirnov (K-S) tests.

## DISCUSSION

Previous work has demonstrated a link between disruptions to the mouse estrous cycle and GnRH neuron function in IHKA mice (Li et al., 2018). Findings from the present study indicate a potential sex-specific role for GABAergic transmission in eliciting changes in GnRH neuron firing activity in this model. In females, these recordings were performed during diestrus, as it was previously observed that increases in GnRH neuron firing were found on diestrus, but not estrus (Li et al., 2018). In the present study, GABAergic transmission to GnRH neurons was increased in diestrous KA-long mice only. This increase was masked when iGluRs were blocked. In males, there was no corresponding increase in the frequency of GABA PSCs, but there was a decrease in PSC half width in the KA group. This finding is indicative of sex-specific responses to the IHKA insult that manifest at the level of GABAergic transmission to GnRH neurons. This study also highlights larger circuit-level considerations when dealing with slice preparations, as traditional pharmacological approaches to isolate GABAergic PSCs impaired circuit integrity to a degree such that a primary phenotypic effect was abolished. In combination with results from previous studies (O’Toole et al., 2014), the present work contributes to the evidence for altered GABA transmission to hypothalamic neuron populations in animal models of epilepsy.

Typically, recordings of fast GABAergic transmission are performed in the presence of iGluR blockers to isolate the GABA_A_R-mediated currents. This is also true for studies of circuits involving GnRH neurons (Adams et al., 2018; Christian and Moenter, 2007; Sullivan et al., 2003). However, after observing that there were no group differences in GABA PSC frequency in the presence of kynurenic acid, we became concerned that the pharmacological iGluR inhibition was disrupting circuit activity in a way that is not representative of the circuit in either control or epilepsy-affected states. Previous work indicated that although sustained glutamatergic transmission can stimulate GnRH neuron action potential firing (Suter, 2004), the simulated level of transmission required exceeds the levels of transmission typically detected perisomatically, even at times of peak GnRH neuron firing activity such as the preovulatory surge (Christian et al., 2009). Together, these results indicate that glutamatergic transmission plays a relatively minor role in regulating GnRH neuron firing activity recorded at the soma, particularly when compared to potent stimulators such as GABA or kisspeptin (Han et al., 2005; Herbison and Moenter, 2011; Pielecka-Fortuna et al., 2008; Zhang et al., 2008).

While the contribution of iGluR-mediated PSCs is relatively small in GnRH neurons (Christian et al., 2009; Liu et al., 2017), this is not the case for important afferent populations, including key GABAergic inputs (Wang et al., 2018). In light of this work, we chose to exclude iGluR-mediated PSCs via a minimum amplitude value cutoff as opposed to circuit-wide pharmacological inhibition, thus allowing us to maintain upstream circuit integrity. Performing recordings in the absence of iGluR blockade, in combination with the cutoff-based approach of excluding iGluR-mediated PSCs, revealed an iGluR-dependent increase in GABAergic transmission to GnRH neurons that was specific to KA-long females. Because this direction of change aligns with previous findings of increased GnRH neuron firing activity in KA-long females during diestrus, it is unlikely that IHKA treatment causes changes in the equilibrium potential of GABA_A_R activation (E_GABA_) such that it would switch from depolarizing (DeFazio et al., 2002) to hyperpolarizing in GnRH neurons. Of note, previous work has highlighted the potential for seizure activity to shift E_GABA_ in a depolarizing direction (Dzhala et al., 2005; Khalilov et al., 2003; MacKenzie and Maguire, 2015; Pathak et al., 2007), including in hypothalamic neurons (O’Toole et al., 2014). However, the findings from this study suggest that a change in the opposite direction may not be as likely, or at least does not occur in GnRH neurons. More specifically, if E_GABA_ is already set at a V_m_ value depolarized relative to the resting membrane potential, it is possible that E_GABA_ is unlikely to shift in a hyperpolarizing direction during states of chronic stress or seizures. Miniature PSCs were not assessed in this study, but it appears that the increased frequency in KA-long mice represents increased activity of presynaptic GABAergic neurons, given that the application of kynurenic acid completely blocked the effect. This finding suggests the presence of an iGluR-dependent increase in feed-forward GABAergic drive. However, increased amplitude of at least some subpopulations of events, both in the presence and absence of kynurenic acid, may indicate changes in the postsynaptic GABA_A_R subunit composition (Verdoorn et al., 1990) and/or changes in quantal size (Auger and Marty, 2000) in IHKA females, both in the presence and absence of comorbid estrous cycle disruption.

In contrast to the findings in IHKA females, there were no observed differences in GABA PSC frequency or amplitude in males. However, there was a decrease in GABA PSC half width in IHKA males that was more prominent than the relatively minor differences in PSC duration in females. This decrease is consistent with stress-induced impairment of GABA_A_R function in rodent models (Biggio et al., 1987, 1981; Drugan et al., 1989). It is unclear whether altered GABA_A_R function plays a role in mediating epilepsy-associated reproductive endocrine dysfunction in males. As such, this potential sex-specific mechanism would be an interesting potential direction of future study.

As the increased transmission occurred exclusively in IHKA females, we can look for potential sources of the altered GABAergic transmission with known sexual dimorphisms in mind. Three major intra-hypothalamic sources of GABAergic transmission to GnRH neurons are the suprachiasmatic (SCN) (Christian and Moenter, 2007; Gu and Simerly, 1997; Van Der Beek et al., 1997), anteroventral periventricular (AVPV), and arcuate (ARC) nuclei (Moore et al., 2015; Yip et al., 2015). Because the kynurenic acid-sensitive increase in GABA transmission was maintained in the coronal slice preparation, the source of the increased GABAergic transmission is likely contained within the slices used for recording. The AVPV has significant overlap with GnRH neuron somata in coronal slices, whereas the locations of the ARC and SCN are too caudal to be maintained in the 300 µm-thick slices used in our experiments (Paxinos and Franklin, 2019). The female-specific finding of increased GABA transmission in IHKA mice also aligns with previous work showing that the AVPV is sexually dimorphic and is larger in females than in males. Within the AVPV, there is a large degree of overlap between GABA and kisspeptin (Cravo et al., 2011). Also of note is the population of ARC kisspeptin neurons, which are glutamatergic and provide an afferent input to AVPV kisspeptin neurons (Wang et al., 2019; Yip et al., 2015). Considered together, we propose a working model in which altered glutamatergic transmission to AVPV kisspeptin neurons (perhaps arising at least in part from ARC kisspeptin neurons) results in a feed-forward mechanism that increases GABAergic transmission to GnRH neurons in diestrous IHKA mice with disrupted estrous cycles (**Fig. 6**).

**Figure 6:**
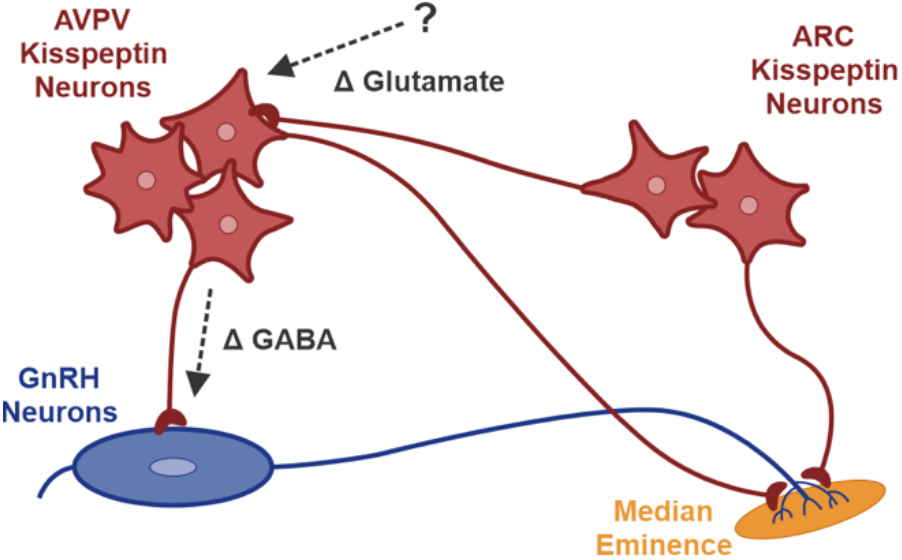
A working model of epilepsy-associated dysfunction in the hypothalamic kisspeptin-GnRH neural circuit in KA-long females. Altered glutamatergic transmission, perhaps arising at least in part from ARC kisspeptin neurons, reaches AVPV kisspeptin neurons, which then release increased amounts of GABA at synapses on GnRH neurons. Note that GnRH neurons send extended projections to the median eminence, which is the site at which GnRH is released into the bloodstream.

### Concluding remarks

The present results provide novel evidence for altered fast synaptic transmission to GnRH neurons as well as their afferent counterparts in IHKA mice. Specifically, these data support a working model in which an epilepsy-associated increase in glutamatergic excitation drives feed-forward depolarizing GABAergic transmission to GnRH neurons, and this phenomenon is specific to females that develop comorbid estrous cycle disruption. These changes in GABAergic transmission also highlight the importance of accounting for sex differences in investigations of epilepsy and its comorbidities (Christian et al., 2020). Considered together, the findings from this study aid in the delineation of the specific pathways and mechanisms linking temporal lobe seizures to dysregulation of hypothalamic circuits controlling reproductive endocrine function.

## ACKNOWLEDGEMENTS

We thank Mia Maren, Jessica Rose, and Cathryn Cutia for assisting with histology experiments.

## FUNDING

This work was supported by the National Institutes Health (NIH)/National Institute of Neurological Disorders and Stroke through grants R01 NS105825 (C.A.C.-H.) and F31 NS124306 (R.J.I.) and by a CURE Epilepsy Research Continuity Fund Grant (C.A.C.-H.).

## AUTHOR CONTRIBUTIONS

Conceptualization: C.A.C.-H.; Formal analysis: R.J.I.; Funding acquisition: C.A.C.-H.; Investigation: R.J.I., L.K.L., V.C.D., J.L.; Methodology: R.J.I., J.L., and C.A.C.-H.; Project administration: C.A.C.-H.; Supervision: C.A.C.-H.; Visualization: R.J.I.; Writing – original draft: R.J.I.; Writing – review & editing: R.J.I. and C.A.C.-H.

## DECLARATION OF INTERESTS

The authors declare no competing financial interests.

## ABBREVIATIONS

ACSF: artificial cerebrospinal fluid
ARC: arcuate nucleus
AVPV: anteroventral periventricular nucleus
GABA: γ-aminobutyric acid
GABA_A_R: GABA_A_ receptor
GFAP: glial fibrillary acidic protein
GnRH: gonadotropin-releasing hormone
iGluR: ionotropic glutamate receptor
IHKA: intrahippocampal kainic acid
KA: kainic acid
LH: luteinizing hormone
MS: medial septum
POA: preoptic area
PSC: postsynaptic current
SCN: suprachiasmatic nucleus
TLE: temporal lobe epilepsy

## Notes

### Competing Interest Statement

The authors have declared no competing interest.

